# Endothelial-targeted CD39 prevents Toxin-induced Pulmonary Hypertension Mice

**DOI:** 10.1101/2025.11.05.686887

**Authors:** Abbey Willcox, Ioanna Savvidou, Natasha Ting Lee, Carly Selan, Ilaria Calvello, Simon C. Robson, Viktoria Bongcaron, Aidan Walsh, Yuyang Song, Xiaowei Wang, Trevor Williams, Karlheinz Peter, Harshal H Nandurkar, Maithili Sashindranath

## Abstract

Disruption of the pulmonary endothelium by drugs, toxins, viruses (e.g., COVID-19), or bacterial sepsis can cause acute pulmonary vasculopathy leading to pulmonary hypertension and consequential heart failure. CD39 is a membrane-anchored ecto-enzyme expressed on endothelial cells (EC), integral in maintaining the antithrombotic profile of the endothelium. This ecto-enzyme works in concert with CD73 to hydrolyze both eATP (pro-inflammatory) and ADP (pro-thrombotic) ultimately to adenosine, which is anti-inflammatory, vasodilatory, and antithrombotic. CD39 activity and adenosine signalling are disrupted in idiopathic pulmonary arterial hypertension (PAH). In this work, we explored the efficacy of endothelial cell-targeted delivery of CD39 to prevent the development of acute toxin-induced PAH in a mouse model.

We generated a novel therapeutic anti-VCAM-CD39 containing an scFv recognising VCAM-1 (a receptor expressed on activated EC) fused to the soluble form of extracellular human CD39. In a mouse model of endothelial cell-toxin-induced PAH, we show that a single administration of anti-VCAM-CD39 (0.4 mg/kg IV) prevented the development of PAH— as reflected in the preservation of right ventricular systolic pressures and the absence of right ventricular hypertrophy at day 10 when compared with controls. This protection is conferred by multiple mechanisms: IL-10-driven potentiation of heme oxygenase (HO)-1, a known inhibitor of smooth muscle proliferation; VCAM-1 blockade reduces leukocyte adhesion to the endothelium; and cytoprotective effects through adenosine signalling. Thus, anti-VCAM-CD39 is a novel bifunctional therapeutic strategy for PAH.

## Introduction

Pulmonary hypertension (PH) encapsulates a group of heterogeneous diseases characterised by an elevation in the mean pulmonary arterial pressures (mPAP) to greater than 20mmHg^1^. Remodeling of the pulmonary vascular bed results in elevation of pulmonary vascular resistance, causing progressive right heart remodeling, hypertrophy, and failure ultimately causing death.^2^ The World Health Organization has standardised PH classification into five groups; in accordance with the underlying aetiology.^3^ These classification efforts differentiate idiopathic pulmonary arterial hypertension (IPAH) (Group 1) from secondary causes of PH. Although these groups display similarities, they are distinct in their aetiologies, pathophysiology, and clinical course. Pre-capillary PH can occur due to lung disease (group 3), chronic thromboembolic disease (group 4), or other rare diseases (group 5). Post-capillary PH results from elevations in mPAPs due to back pressure from the pulmonary venous system or left side of the heart (group 2). Idiopathic PAH (group 1) is a rare condition characterised by pre-capillary PH without other causes.

PH remains an incurable disease with a 43% five-year mortality despite modern treatment options^4^ underscoring the need for innovative translational strategies to target novel pathways involved in onset and progression. Several acute insults such as sepsis or viral infections are linked to PH^5–7^, though hypoxia or shear stress^8^ induced endothelial damage are likely to be implicated. The pathogenic role of endothelial dysfunction has been increasingly appreciated with imbalances in vasoactive endothelial mediators such as prostacyclin, nitric oxide (NO), and endothelin-1.^4,9^ Inflammation plays an integral role in the initiation and progression of PAH^10,11^. Endothelial cells from patients with PH express an abnormal inflammatory phenotype, characterised by surface expression of E-selectin, ICAM-1, and VCAM-1^11^ as well as heightened levels of inflammatory cytokines such as TNF-α and IL-6, which appears to correlate with poorer patient outcomes^12,13^. One of the hallmarks of inflammation is increased tissue metabolism, which invariably increases local oxygen demand and results in tissue hypoxia. Emerging data suggests that Hypoxia-inducible factor (HIF) signalling plays a critical role in the initiation and progression of all PH categories but specifically PAH.^14–16^ In hypoxic conditions, HIF-⍺ subunits become stabilised, which allows for the HIF-⍺ and HIF-β subunits to bind, initiating adaptive responses to hypoxia by modulating vascular tone, angiogenesis, erythropoiesis, cell proliferation, and survival, along with autophagic responses.^17^ Under the conditions of hypoxic stress, important transcription regulators of pulmonary artery smooth muscle cell (PASMC) proliferation are HIF1-α as a stimulator and haem oxygenase-1 (HO-1) as an inhibitor^15,16,18^. Elevated HIF1-⍺ has been described in both primary and secondary PAH,^19–24^ where the principal source is thought to be pulmonary arterial endothelial cells and PASMCs.^24–26^ Interleukin (IL)-10 is a potent anti-inflammatory cytokine. In PAH, IL-10 inhibits TGF-β, a key promoter of PASMC proliferation, upregulates HO-1, and induces the expression of CD39.^27^ ^18,28,29^

Expression and enzymatic activities of CD39 and CD73 regulate extracellular ATP/adenosine ratio, which influences pulmonary vascular homeostasis and potentially plays a central role in the pathophysiology of PH.^30^ An ATP-rich environment has been shown to promote pulmonary smooth muscle migration and proliferation.^30^ CD39 expression is downregulated on pulmonary ECs in patients with PAH. ^30,31^ Concordantly, mice with targeted deletion of CD39 have a higher ATP:adenosine ratio and develop PH especially under conditions of hypoxia^30,31^ . Owing to its vasodilatory activity, adenosine has long been of interest as a therapeutic strategy in PAH^32^. Adenosine infusion resulted in reduction in right ventricular pressure in newborn lambs with hypoxia-induced PH^33^; however, response to adenosine in patients appears to be dependent on the underlying cause of the PAH, with only 10–20% of patients with IPAH exhibiting a reduction in their PAP with adenosine infusion.^34,35^ Systemic delivery of adenosine as a therapeutic strategy is hampered by its exceptionally short half-life (5–10 seconds) and its propensity for systemic side effects including bradycardia and hypotension.^36^ Nevertheless, the potent roles of purinergic nucleotides suggest that novel ways of altering the ATP:adenosine ratio selectively in the microvasculature and thereby abrogating the need for systemic infusion of adenosine are an appealing therapeutic strategy. Targeted strategies for delivering CD39 have been promising in preclinical models of cardiac ischaemic reperfusion injury^37^ and sepsis^38^ where CD39 was targeted to activated platelets.

We developed a novel therapeutic, anti-VCAM-CD39, comprised of a single-chain antibody recognising VCAM-1, a receptor expressed on activated endothelial cells, fused to human soluble CD39. We have shown previously that anti-VCAM-CD39 has the bifunctional effect of blocking the trans-endothelial migration of migration, comparable to a commercially available neutralising VCAM-1 antibody whilst also enhancing adenosine generation. These effects confer protection in a mouse model of unilateral stroke ^39^ and in global fore brain hypoxic injury^40^ . We demonstrate here in a proof-of-concept murine model that the endothelial-targeted bifunctional activities of our novel therapeutic anti-VCAM-CD39, will prevent the development of PAH after monocrotaline pyrrole (MCTP)-induced endothelial injury.

## Methods

### Design of the anti-VCAM-CD39 Expression Construct and Protein Production

Design and production of anti-VCAM-CD39 has been previously described in prior publications^40,41^ .

### Human lung tissue

De-identified human specimens were collected with the assistance of the Alfred Hospital Pathology Department and approved by the Alfred Hospital Ethics Committee (Project number 158/21, April 2021).

### Mice

All animal protocols were reviewed and approved by the Alfred Research Alliance Ethics Committee (approval number E/8254/2022/M). Adult male BALB/c mice (seven weeks of age, 28–30 g) were purchased from a breeding colony at Walter & Eliza Hall Institute for Medical Research.

### PAH Model

PAH was induced in 7-week-old male Balb/C mice using a single IV injection of the endothelial toxin MCTP 8 mg/kg (Santa Cruz, A3118, diluted in 2% DMF in RPMI to a final concentration of 2.5 mg/ml) as previously described.^42,43^.

Pulmonary pressures were periodically assessed by functional rodent echocardiography^44^ and right heart catheterisation 10 days after MCTP injection. Pulse-wave doppler echo was used to record the pulmonary blood outflow at the level of the aortic valve in the short-axis view to measure the pulmonary acceleration time (PAT) and pulmonary ejection time (PET). The PAT/PET ratio is inversely correlated to right ventricular pressures^44^ . See Supplementary Material for further information.

### Treatment Administration

Anti-VCAM-CD39 (0.4 mg/kg IV) was injected 72 hrs post MCTP administration. Equivalent doses of the controls were administered; non-targeted CD39 which contained an scrambled non-binding VCAM (ScrVCAM-CD39, 0.4mg/kg) and VCAM antibody alone (anti-VCAM 0.13 mg/kg). All agents were diluted in TBS and given via IV tail vein injection.

### Euthanasia and Tissue Harvesting

At D10 post-MCTP injection, mice were anaesthetized with pentobarbitone Sodium (90 mg/kg, Provet Pty Ltd.), and transcardially perfused with phosphate-buffered saline. The RV was dissected, and the Fulton’s Index was calculated:

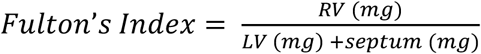

Lungs were harvested; the left lobe was inflated with Tissue-Tek OCT and snap-frozen in chilled isopentane on dry ice. The remaining right lung was homogenised in Lysis Buffer (TBS +1% Triton X-100; Roche) or RNAlater (Thermo Fisher Scientific, Australia) at 300mg wet tissue per 1 ml.

### Histology

OCT-embedded lung tissue was cryosectioned at 6µm and stored at –80 ℃.

#### Haematoxylin and Eosin (H&E) and Carstairs stain

Cryosections (6 µm) were stained with H&E by Monash Histology Platform using standard protocols as described^45^.

#### Smooth muscle actin (SMA) Immunofluorescence

Refer to Supplementary Material for further information

### Immunofluorescent Staining

Refer to Supplementary Material for further information

### RT-PCR

RT-PCR was performed using the SensiFAST Lo-ROX Probe Kit (Bioline, Australia). Expression of ICAM1, IL-6, IL-1β, TNF⍺, HIF1⍺, IL-10, HO-1, and TGFβ was assessed, with gene expression assays purchased from Thermo Fisher Scientific, Australia. RT-PCR was run on the QuantStudio 6 Real-Time PCR System (Thermo Fisher Scientific, Australia) using the standard program. Refer to Supplementary Material for further information

### eNOS ELISA

Refer to Supplementary Material for further information

### Statistical Analysis

Statistical analysis was performed using Prism 9 software (GraphPad, US). Normality tests were run to determine subsequent statistical test. Confirming normality, comparison of experimental datasets was performed by one-way ANOVA with Dunnett’s post-hoc correction or two-way ANOVA with Sidak’s or Dunnett’s post-hoc correction as stated. Non-normal datasets were compared with Kruskal-Wallis test with Dunn’s multiple comparisons test. Differences between two groups were assessed by two-tailed student t-tests (unpaired or unpaired with Welch’s correction for parametric data and Mann-Whitney test for non-parametric data). P< 0.05 was considered significant.

### Study Approval

All experiments were approved by Alfred Research Alliance Ethics Committee (ethics applications number E/8254/2022/M) in accordance with Australian code for the care and use of animals for scientific purposes 8th Ed 2013 and in compliance with ARRIVE guidelines for reporting animal experiments.

### Data availability

Values for all data points in graphs are reported in the Supporting Data Values file. Any additional data can be requested from corresponding author AW.

## Results

### PAH is associated with reduced expression of CD39 and presence of endothelial activation with upregulation of VCAM-1

Lung tissue from patients was assessed for disruption of purinergic markers and indicators endothelial activation compared with a control group (non-PAH patients). Figure 1a shows representative histology from a patient with PAH, demonstrating reduction in the CD39 expression in the pulmonary vessels immunostained for CD31, compared with the non-PAH patient. Furthermore, endothelial dysfunction can be seen in the lung tissue of patients with PAH as demonstrated in Figure 1b, where representative images show upregulation of VCAM-1 (green) on pulmonary vessels from a patient with PAH compared to a patient without PAH.

**Figure 1:**
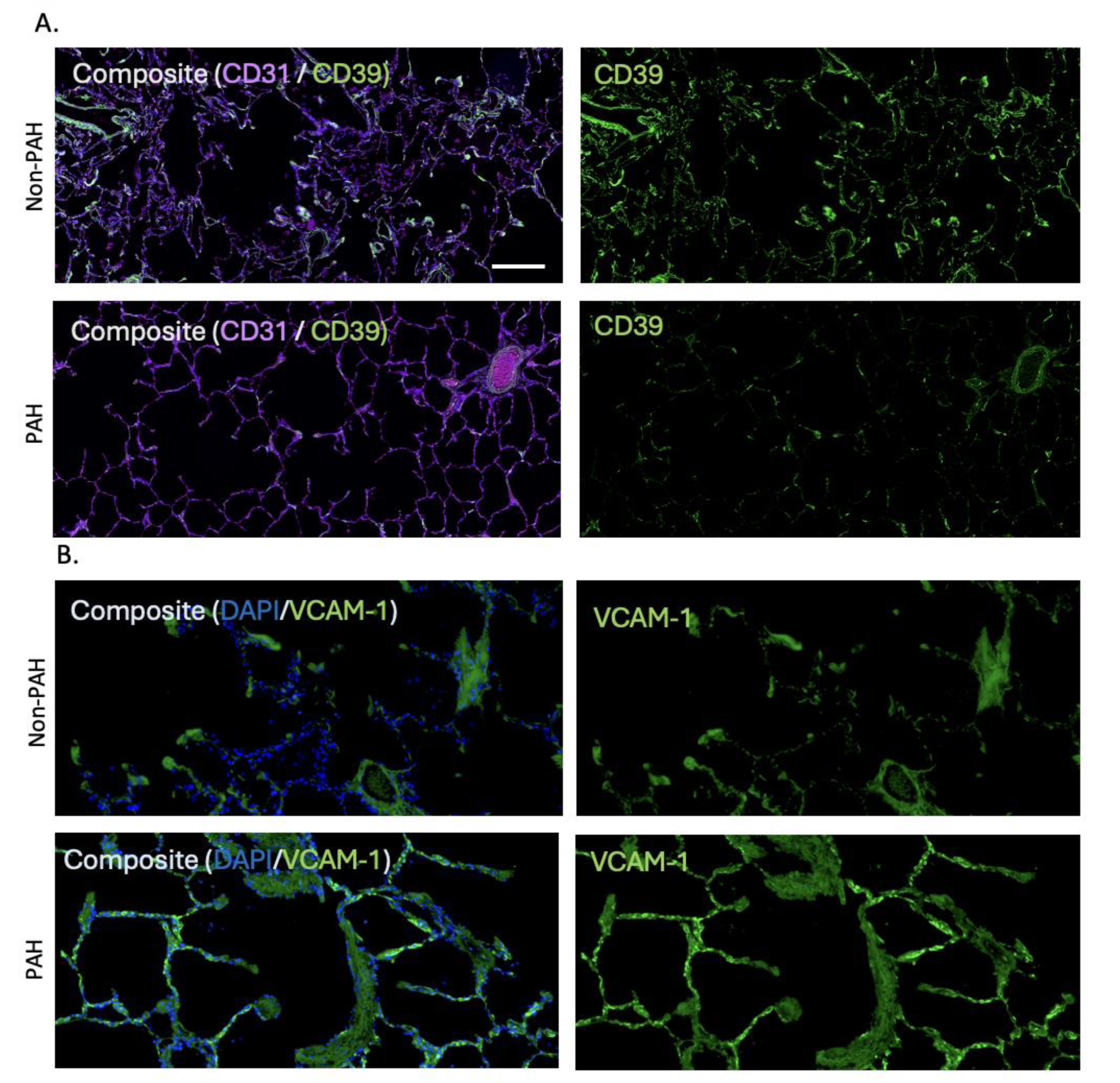
CD39 Immunoreactivity is diminished in the lungs of patients affected by PAH and VCAM is upregulate; A) Representative immunofluorescent staining of lung tissue from a patient with PAH (second panel) demonstrates significant reduction in CD39 (green) staining on the endothelial vessel (CD31, purple) compared with the control (top panel). B) Representative immunofluorescent staining of lung tissue from a patient with PAH (second panel) demonstrates increase in VCAM-1 staining (green) compared with the non-pulmonary hypertension patient (top panel). Blue staining is DAPI.

### Monocrotaline pyrrole promotes endothelial activation and inflammation in the pulmonary vasculature and the development of PAH in Mice

Male BALB/c mice (age seven weeks) were injected with a single IV dose of MCTP (8 mg/kg) or VC (DMF 2%) at day 0 (D0) to induce PAH.

Using synchronous echocardiogram and right heart catheterisation an inverse correlation between the PAT/PET ratio and the RV pressures was observed (Fig 2A, Pearson r = 0.49, p = 0.021). MCTP-treated animals had a significantly shorter PAT/PET ratio than VC-treated animals (Fig 2B; median ratio for VC 0.45 versus for MCTP 0.30, p < 0.001) at D10. The reduction in PAT is reflected in a distinctly different flow curve (Fig 2C). MCTP treatment was associated with elevated right ventricular pressures (RVSP) by D10 (Fig 2D, median pressure for VC 27.5mmHg versus for MCTP 39.8mmHg, p = 0.003). Quantification of the Fulton’s index confirmed that elevated RVSP correlated with the development of right ventricular hypertrophy (Fig 2E, VC 0.230 versus MCTP 0.293, p = 0.0027). Exposure to MCTP resulted in vessel wall remodeling with arteriolar medial thickening appreciable on the H&E stain (Fig 2F). There was evidence of endothelial activation with VCAM-1 upregulation on immunofluorescence correlating with CD31 expression (Fig 2G).

**Figure 2.**
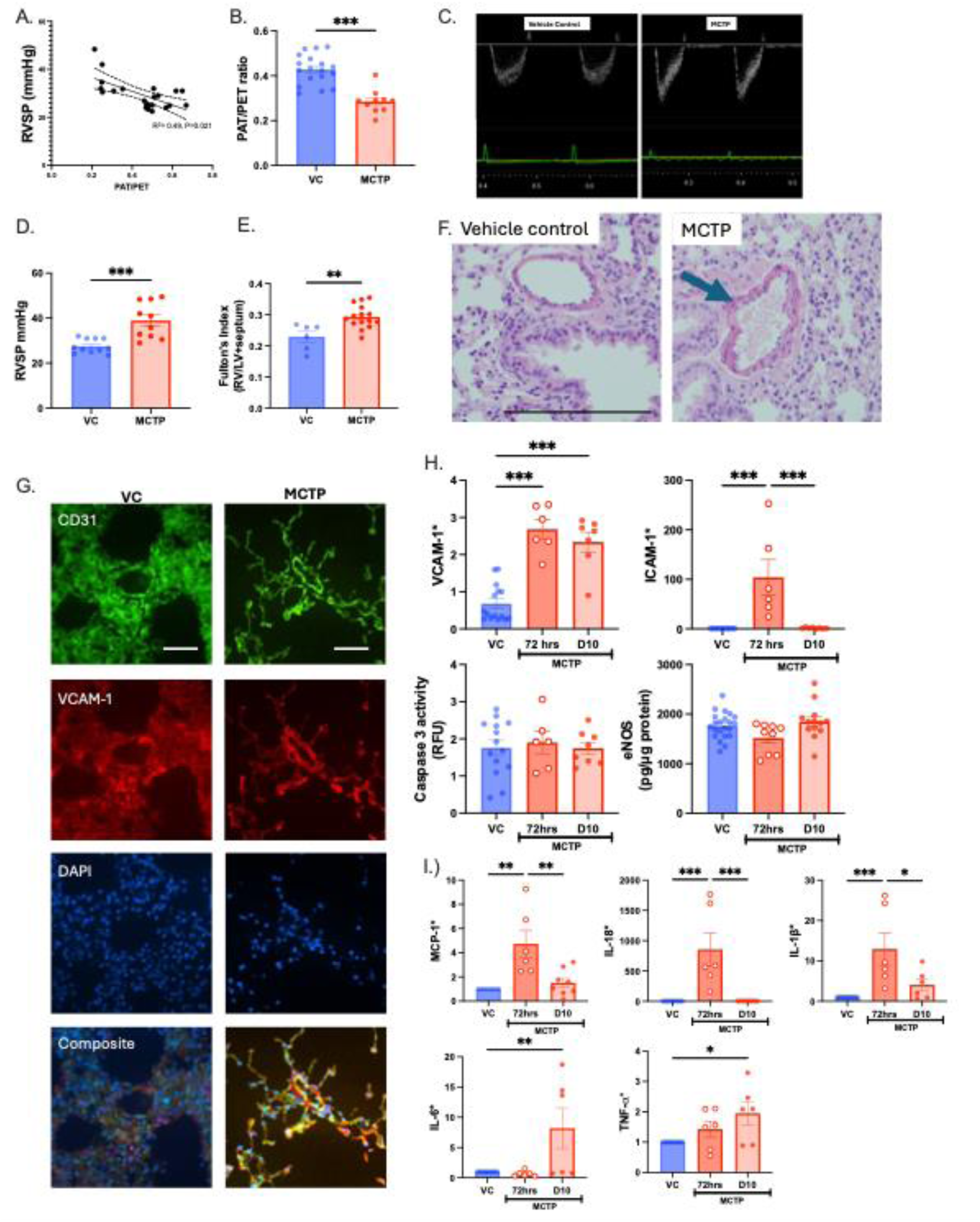
MCTP induces PH in mice at 10 days after injection. Non-invasive echocardiogram monitoring correlates inversely with RVSP (A), with a decline in the PAT/PET ratio as the RVSP rises. The PAT/PET ratio was reduced at D10(B). Flow curves over the pulmonary valve change dramatically, representative image of D10 (C). MCTP (red) induces elevated RVSP (D.) and right ventricular hypertrophy measured as the Fulton’s Index (E) within 10 days of administration compared with VC (blue) consistent with the development of toxin-induced PH complicated by right ventricular hypertrophy. Pulmonary vascular remodeling with medial wall hypertrophy on a representative H&E stain (F) with associated VCAM-1 upregulation (G) is shown at D10 post-injection. Pulmonary endothelial activation is observed within 72 hours of MCTP injection with rise in VCAM-1, ICAM-1 without significant change in Caspase 3 and eNOS (G). Acute inflammation is observed with increase in IL-18, MCP-1 and IL-1β at 72 hrs and IL-6 and TNF⍺ at D10. Data are expressed as mean ± SEM, n = 6 or more per group. * H and I data expressed as fold change relative to VC unless otherwise specified. Unpaired Welch’s T test (B, D, E) or Ordinary one-way ANOVA with Tukey’s multiple comparisons test ( H, I). *p < 0.05, **p < 0.01, ***p < 0.001. Black scale bar = 200um, white scale bar = 50um MCTP (monocrotaline pyrrole), RVSP (right ventricular systolic pressure), VC (vehicle control), PAT (pulmonary acceleration time), PET (pulmonary ejection time), SMA (smooth muscle actin).

Maximal upregulation of both VCAM-1 (VC 1.22 versus MCTP 2.56, p = 0.0085) and ICAM-1 (VC 1.0 versus MCTP 104.2, p = 0.001) was observed in the lungs 72 hours after MCTP injection. There was differential time course of expression for VCAM-1 and ICAM-1. While VCAM-1 remained significantly higher at D10, ICAM-1 expression had returned to the baseline (Fig 2H), justifying our strategy of using VCAM-1 as the endothelial epitope for the targeted delivery of CD39.

MCTP-induced PAH is preceded by early inflammatory changes in the lung with elevation in IL-18, MCP-1, and IL-1β within 72 h of injection while expression of IL-6 and TNF-⍺ was elevated at day 10 post injection (Fig 2I).

### Anti-VCAM-CD39 abrogates MCTP-induced PAH in a mouse model

MCTP-injected animals were treated at 72 h, the point of maximal VCAM-1 upregulation, with anti-VCAM-CD39 or control, and assessed for evidence of PAH at D10. Administration of anti-VCAM-CD39 completely ameliorated the development of PAH with normal RVSP on D10 (Fig 3A) and preservation of the PAT/PET ratio (Fig 3B). Most importantly, RV hypertrophy was not observed in animals treated with anti-VCAM-CD39 (Fig 3C). Animals treated with anti-VCAM appeared to have normal measured RVSP by D10 along with higher PAT/PET ratios compared to TBS-treated animals suggestive of protection, however, these animals had evidence of RV remodeling and hypertrophy (Fig 2C). In contrast, untargeted CD39 (ScrVCAM-CD39) did not protect animals from developing PAH with persistent reduction in PAT/PET ratio, raised RVSP and RV remodeling and hypertrophy. These data confirm the bifunctional activity of anti-VCAM-CD39 with contributions from both VCAM-blockade and CD39 enzyme activity as well the advantage of endothelial targeting of CD39 as equimolar quantity of untargeted CD39 (scrVCAM-CD39) had no benefit.

**Figure 3:**
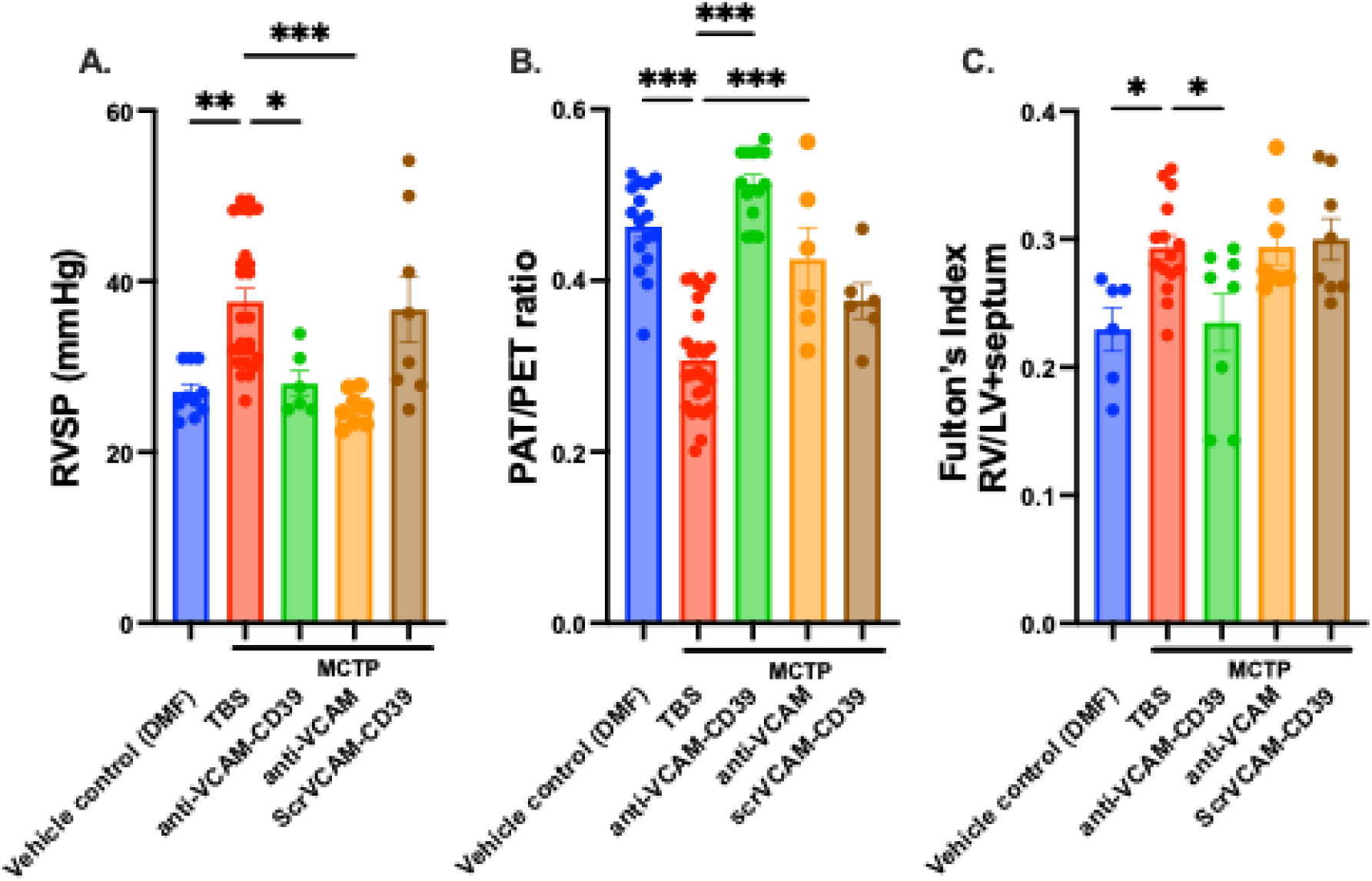
Anti-VCAM-CD39 abrogates the development of MCTP-induced PAH. Animals treated with the VC or MCTP IV (8 mg/kg) followed by treatment at 72 hours with TBS, anti-VCAM-CD39, anti-VCAM or scr-VCAM-CD39. A: At D10, RVSP is normal in animals treated with anti-VCAM-CD39 and anti-VCAM compared with those treated with TBS or scr-VCAM-CD39. B: The PAT/PET ratio is normal in animals treated with anti-VCAM-CD39 and, to a lesser extent, with anti-VCAM and scr-VCAM-CD39, compared with animals treated with TBS. C: Right heart hypertrophy is measured using the Fulton’s Index (RV/LV + septum). Anti-VCAM-CD39-treated animals have preserved RVs, whereas groups treated with either TBS, anti-VCAM or scr-VCAM-CD39 have evidence of right ventricular hypertrophy with elevated Fulton’s Index. MCTP = monocrotaline pyrrole, TBS = tris buffered saline, RVSP = right ventricular systolic pressure, PAT = pulmonary acceleration time, PET = pulmonary ejection time, RV = right ventricle, LV = left ventricle, PAH; pulmonary arterial hypertension, DMF; dimethylformamide, VCAM; vascular adhesion molecule-1. Data were analysed with one-way ANOVA analysis using Tukey’s multiple-comparison post-hoc test (Prism 9). Data are mean ± SEM unless otherwise specified, n > 6. ns; p > 0.05, *p < 0.05, **p < 0.01, ***p < 0.001.

### Anti-VCAM-CD39 abrogates MCTP-Induced pulmonary vessel remodeling

MCTP induced lung vessel remodeling with fibroelastic intimal hyperplasia and concentric medial hyperplasia notable on the H&E staining (Fig 4A). Carstairs staining highlights fibrin strands throughout the vessels (Fig 4A, indicated by the red ‘arrows’) in the MCTP-treated animals. This vessel remodeling and fibrin deposition were not present after treatment with anti-VCAM-CD39. The reduced vascular remodeling was further corroborated by the assessment of SMA staining via immunofluorescence. Exposure to MCTP resulted in significantly more SMA deposition on D10 which was prevented with anti-VCAM-CD39 treatment (Fig 4B). Such attenuation of MCTP-induced vessel remodeling was not seen with the anti-VCAM alone nor untargeted CD39 (Fig 4B), thereby confirming the combined advantage of VCAM-targeting and endothelial enrichment of CD39 enzyme activity.

**Figure 4:**
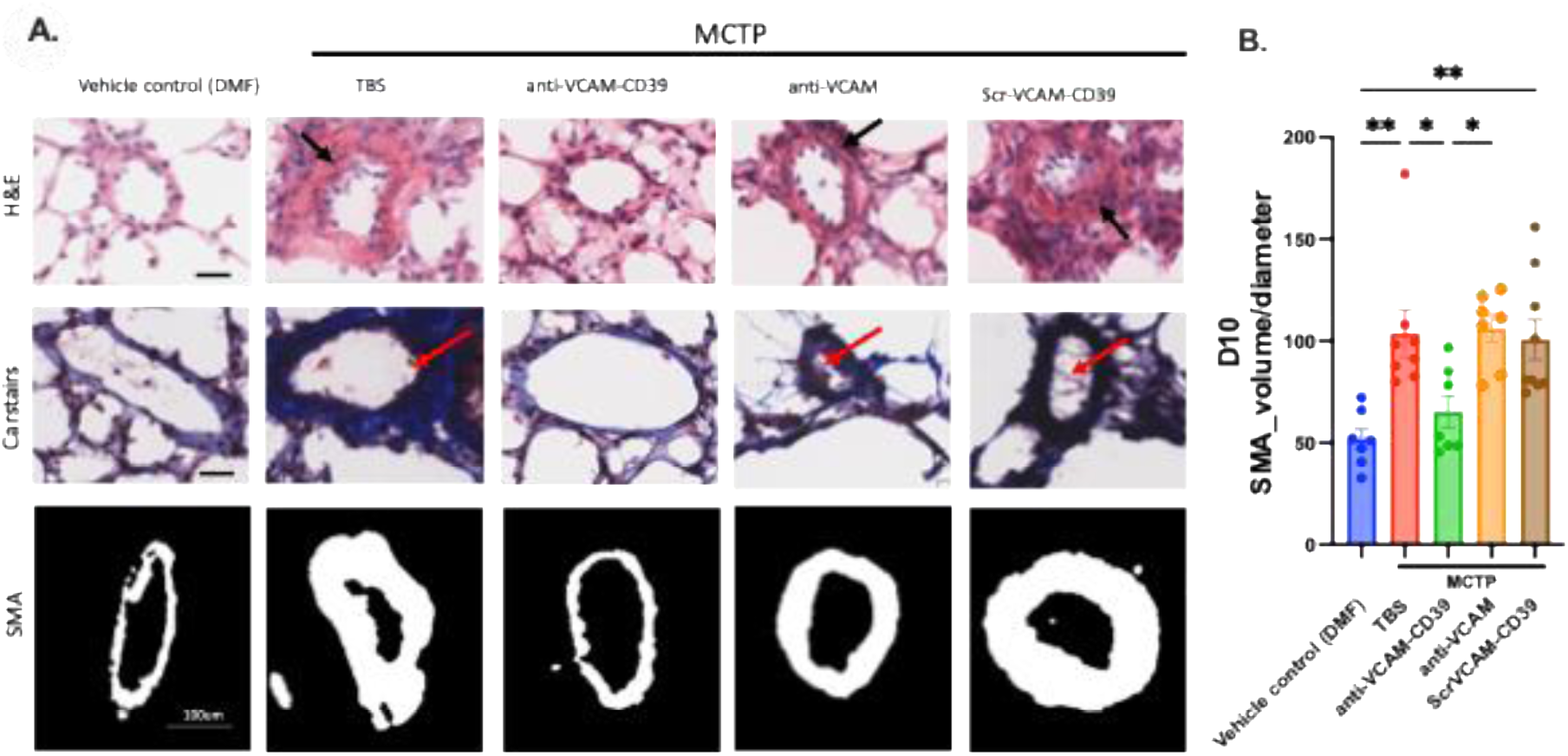
Anti-VCAM-CD39 prevents lung blood vessel remodeling in MCTP-induced PAH in a mouse. A) Representative histology of D10 lung sections from animals treated with the VC (DMF) or MCTP 8 mg/kg followed by TBS, anti-VCAM-CD39, anti-VCAM or scr-VCAM-CD39. Significant remodeling with concentric wall thickening from muscle proliferation and intimal hypertrophy (short black arrow) can be seen in animals treated with TBS, anti-VCAM and scr-VCAM-CD39. Mild pulmonary vessel wall thickening can be seen in the anti-VCAM-treated animals. Very little to no vascular remodeling is noted in the anti-VCAM-CD39-treated animals compared with control animals. Fibrin strands are apparent on Carstairs stain in animals treated with scr-VCAM-CD39, TBS and anti-VCAM (red arrow). Representative histology of D10 lung sections stained for SMA (A) and quantification of SMA (B) demonstrate significant SMA deposition in the small vessels of animals treated with MCTP followed by TBS, scr-VCAM-CD39 and anti-VCAM, but not after treatment with ant-VCAM-CD39.. Data are mean ± SEM, n = 6 for each group. *p < 0.05, **p < 0.01. Black scale bar = 25um.

### Anti-VCAM-CD39 reduces inflammation caused by MCTP-Induced PAH

MCTP-induced PAH exhibits a pro-inflammatory phenotype with elevations of IL-6 and IL-1β and TNF-⍺ expression (Fig 5) on D10. We observed that anti-VCAM-CD39 treatment had the most effect in significantly suppressing the gene expression IL-6 and IL-1β (Fig 5A & B). TNF-⍺ expression however remained unchanged by anti-VCAM-CD39 treatment (Fig 5C).

**Figure 5:**
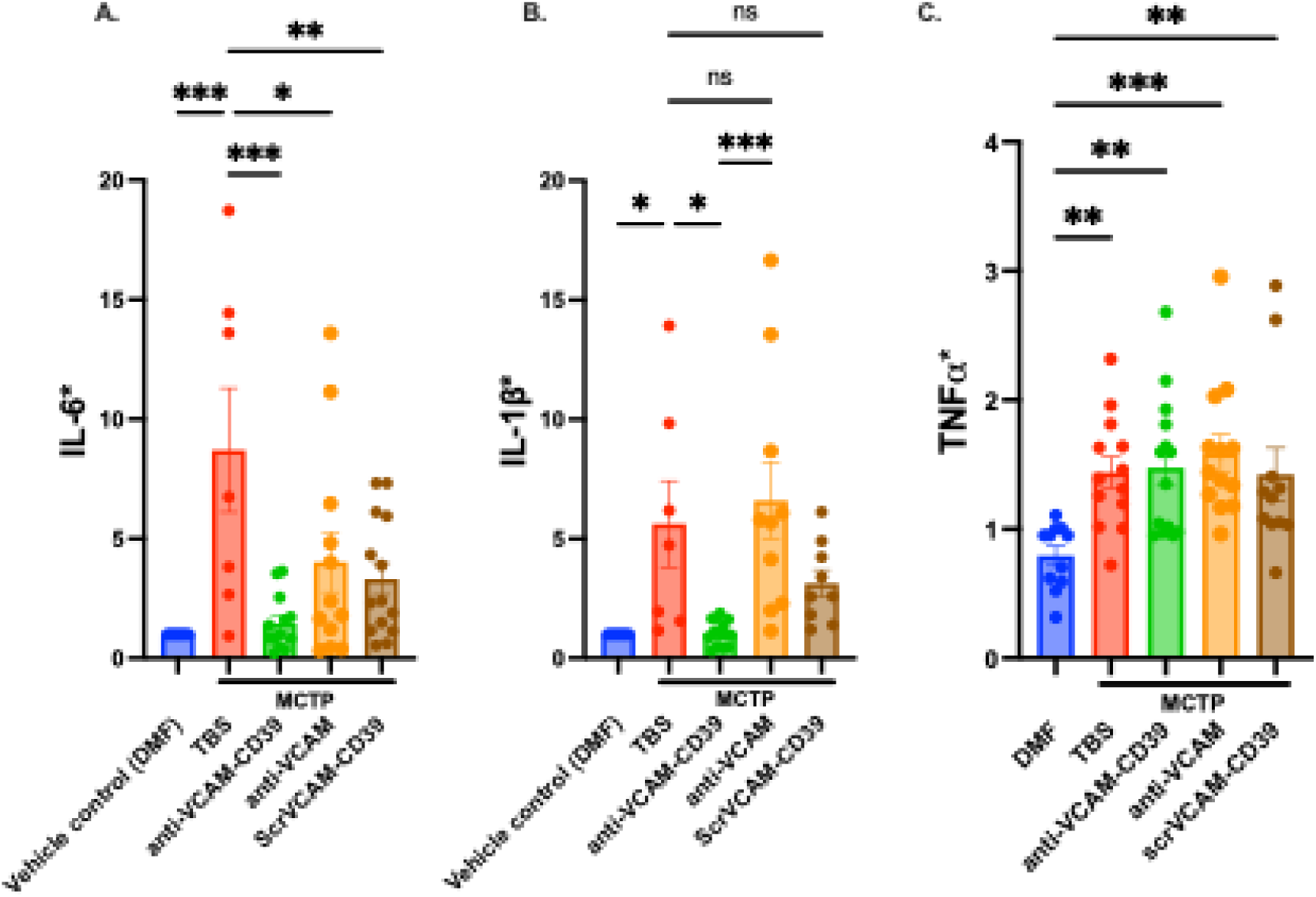
Treatment with anti-VCAM-CD39 reduces MCTP-induced inflammation at D10 The MCTP-induces a rise in expression of inflammatory cytokines. Anti-VCAM-CD39 treatment abrogates this. In IL-6 and IL-1β (A & B) compared with other controls at D10 but not the rise in TNF-⍺ (C). *Data are expressed as fold change from VC-treated animals (DMF). Data are mean ± SEM, n = 6 for each group, p > 0.05, *p < 0.05, **p < 0.01, ***p < 0.001.

### Endothelial targeted CD39 likely protects from MCTP-Induced PAH development through IL-10-mediated HO-1 inhibition of PASMC proliferation and correction of hypoxia

MCTP-induced endothelial injury and consequent changes in intrapulmonary architecture result in hypoxic stress marked by the elevation in the transcription of HIF1-⍺. The bifunctional actions of anti-VCAM-CD39 are necessary to ameliorate hypoxia with reduction in HIF1-⍺ (Fig 6A). Additional contribution is made by elevations in the levels of the anti-inflammatory cytokine IL-10 only in the cohort treated with anti-VCAM-CD39 (Fig 6B). IL-10 can have the dual actions of increasing HO-1 activity and inhibiting TGF-β to regulate pulmonary smooth muscle cell proliferation. In our model of PAH, we observed increment in HO-1 solely with anti-VCAM-CD39 (Fig 6C) but no changes in TGF-β (Fig 6D).

**Figure 6:**
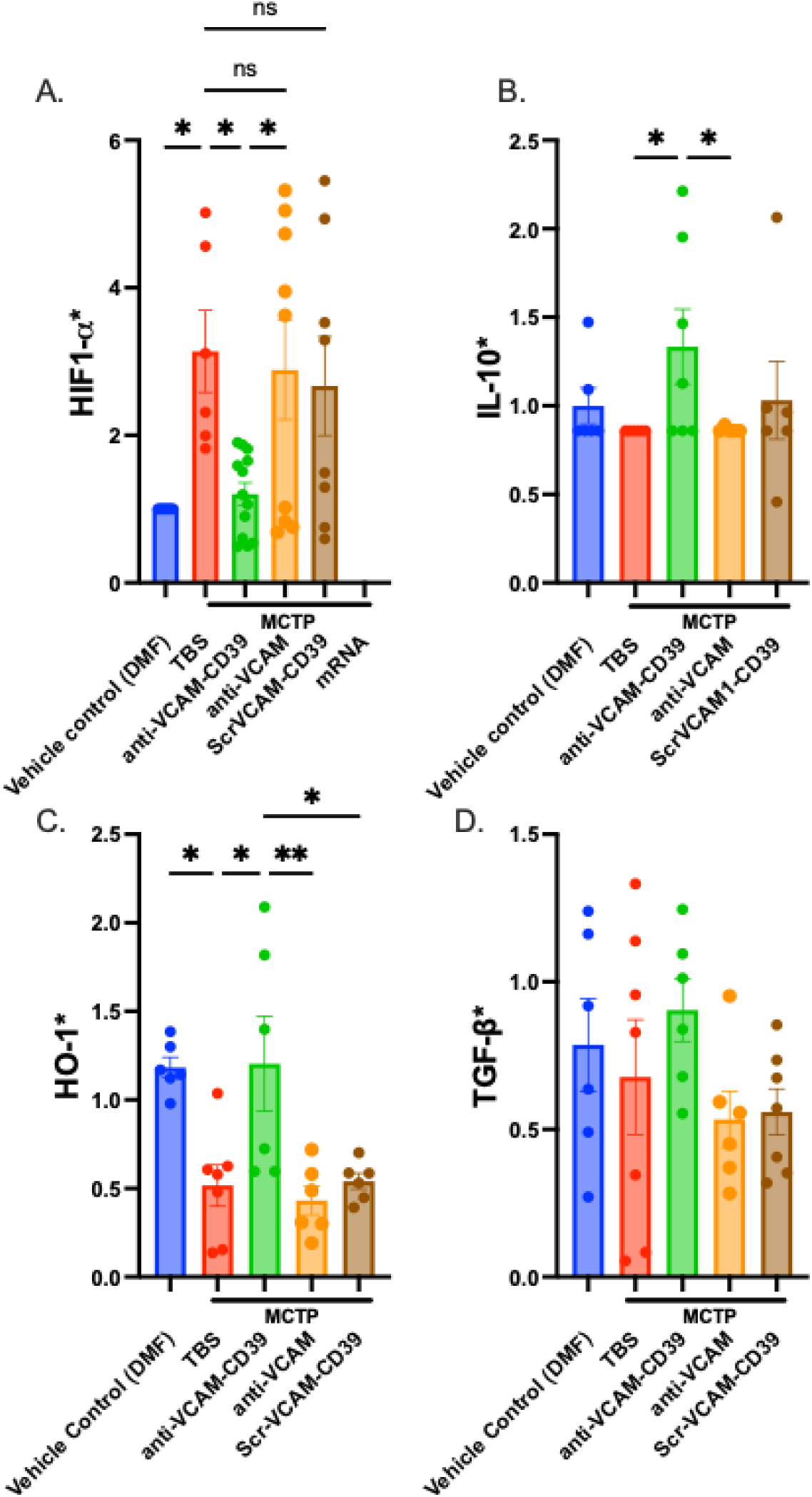
Anti-VCAM-CD39 pulmonary oxygenation and reduces pulmonary vascular remodeling via potentiation of IL-10 driven HO-1. Reduction in the transcription of HIF1-⍺ in the lungs by Anti-VCAM-CD39 demonstrates amelioration of tissue hypoxia (A). IL-10 expression is increased significantly in animals treated with anti-VCAM-CD39(B). Expression of HO-1 is restored in MCTP-treated animals who received anti-VCAM-CD39 compared with those who received TBS, anti-VCAM, or ScrVCAM-CD39 (C) without any change in TGF-β expression(D). *Data are expressed as fold change from VC-treated animals (DMF). Data are mean ± SEM, n = 6 for each group, p > 0.05, *p < 0.05, **p < 0.01, ***p < 0.001

## Discussion

Pulmonary hypertension is characterised by a pro-inflammatory signature of activated vascular endothelial cells and disrupted purinergic signalling^32^ . Cultured ECs from patients with PAH appear to have an abnormally activated phenotype characterised by surface expression of E-selectin, ICAM-1, and VCAM-1.^8,11^ Similar to prior studies^11,46^, we demonstrate upregulation of VCAM-1 in both patients with PAH and in mice with toxin-induced PAH. In addition to being a marker of endothelial activation, VCAM-1 facilitates adherence and transmigration of leukocytes thereby contributing to a proinflammatory phenotype. We have previously demonstrated that the binding of anti-VCAM-CD39 to VCAM-1 inhibits leukocyte transmigration comparable with the effect of a VCAM-1 specific monoclonal antibody^47^ . This action contributes to the therapeutic benefit of Anti-VCAM-CD39. However, as anti-VCAM alone is less effective than anti-VCAM-CD39, it proves the importance of CD39 activity.

Visovatti et al.^30^ described a severe phenotype of hypoxia-induced PAH in CD39-knockout mice which could be salvaged using soluble apyrase (an endonucleotidase derived from potato extract with ATPase and ADPase activity like CD39). In addition, apelin, an endogenous peptide known to potentiate CD39,^31^ can ameliorate the deleterious effects of PH in both animals and patients.^48,49^ Although soluble CD39 is antithrombotic, dose-dependent prolongation of bleeding times has been consistently shown^50–52^. We have taken advantage of VCAM-1 upregulation by using this receptor to target the enzymatic activity of CD39 specifically to the endothelial surface so that it can be administered at a dose of 0.4mg/kg which is considerably lower than the dose ≥ 2mg/kg-that increases bleeding^39^. We have already demonstrated the efficacy of this strategy with use of anti-VCAM-CD39 in unilateral stroke ^39,47^ and global brain hypoxia^40^ . Using a similar approach, low dose CD39 targeted to activated platelets ameliorates cardiac ischaemic reperfusion injury^37,53^ . This advantage also explains why equimolar doses of non-targeted recombinant CD39 (scrVCAM-CD39) is less effective without the benefit of concentration on activated endothelium. The temporal profile of endothelial activation correlated with disease onset in this animal model and provided invaluable rationale for the timing of anti-VCAM-CD39 delivery in this model.

Consistent with our findings, purinergic signalling dysfunction, notably loss of pulmonary CD39, has been previously described in patients and animal models of PAH,^30,31^ with some suggesting that loss of CD39 correlates with severity of vessel remodeling in PAH.^31^ Furthermore, cultured PAECs lacking CD39 appear to have an apoptosis-resistant phenotype,^31^ which is also a feature in PAECs from patients with PAH.^54^ Given the importance of CD39 in scavenging eATP and promoting adenosine, the loss of CD39 may ostensibly be the underlying cause of the adenosine-poor state seen in patients with PAH.^55^ and justifies our strategy for CD39 enrichment.

HIF1-⍺ is known for its adaptive response mechanism in lung injury. We observed that MCTP-induced HIF1-⍺ expression was significantly reduced in animals who received anti-VCAM-CD39 compared with those who received control treatments. The role of HIF1-⍺ has been controversial,^14^ with some studies showing that deletion (or partial deletion) of HIF1-⍺ protected mice from developing PH and/or RV hypertrophy after chronic hypobaric hypoxia.^56,57^ However, Ball et al.^58^ reported that inducible SMC-specific deletion of HIF1-⍺ attenuated PH but did not affect RV hypertrophy, and another study showed that mice with constitutive SMC-specific HIF1-⍺ deletion exacerbated hypoxia-induced PH.^59^ More recent studies have demonstrated that mice with constitutive EC-specific HIF1-⍺ deletion are not protected from hypoxia-induced PH and RV hypertrophy.^60,61^ Although chronic hypoxia is known to be the most common way in which HIF are activated, its involvement in various sub-types of PH suggests other factors such as endothelial dysfunction, inflammation or vasoconstriction may be able to activate HIFs.^14,17,62^

Pro-inflammatory cytokines such as TNF-⍺ and IL-1β have been shown to stabilise HIF1-⍺ via NF-κB.^63^. Consistent with this, we observed that the elevated IL-1β expression seen in the TBS and anti-VCAM-treated animals which mirrored with elevated HIF1-⍺ expression. The lower HIF1-⍺ seen in the anti-VCAM-CD39-treated animals may be reflective of reduced hypoxia and tissue inflammation which results in less smooth muscle proliferation and prevents the development of PAH.^64^ Previous studies have shown that induced IL-10 expression prevented the development of MCT-induced PAH in rats^64^ and enhancement of HO-1 in the lung prevented the development of hypoxia-induced PAH and notably inhibited the structural remodeling of the pulmonary vessels^65^. The role of adenosine in augmenting IL-10 expression,^66–68^ as well as the effects of HO-1 in reducing acute pulmonary inflammation via adenosine receptors,^69^ have been well characterised. We have previously shown that anti-VCAM-CD39 shifts the ATP: adenosine ratio in favour of an adenosine-rich environment^39^. Together this suggests that the likely mechanism of the protective effects of anti-VCAM-CD39 on pulmonary vascular remodeling and subsequent pulmonary hypertension is via localised and targeted adenosine-mediated increase in expression of IL-10, which reduces vascular remodeling via promotion of HO-1 (Fig 7). It remains to be determined whether the concomitant reduction in eATP at the pulmonary endothelial surface provides an alternative or additive mechanism of protection.

**Figure 7:**
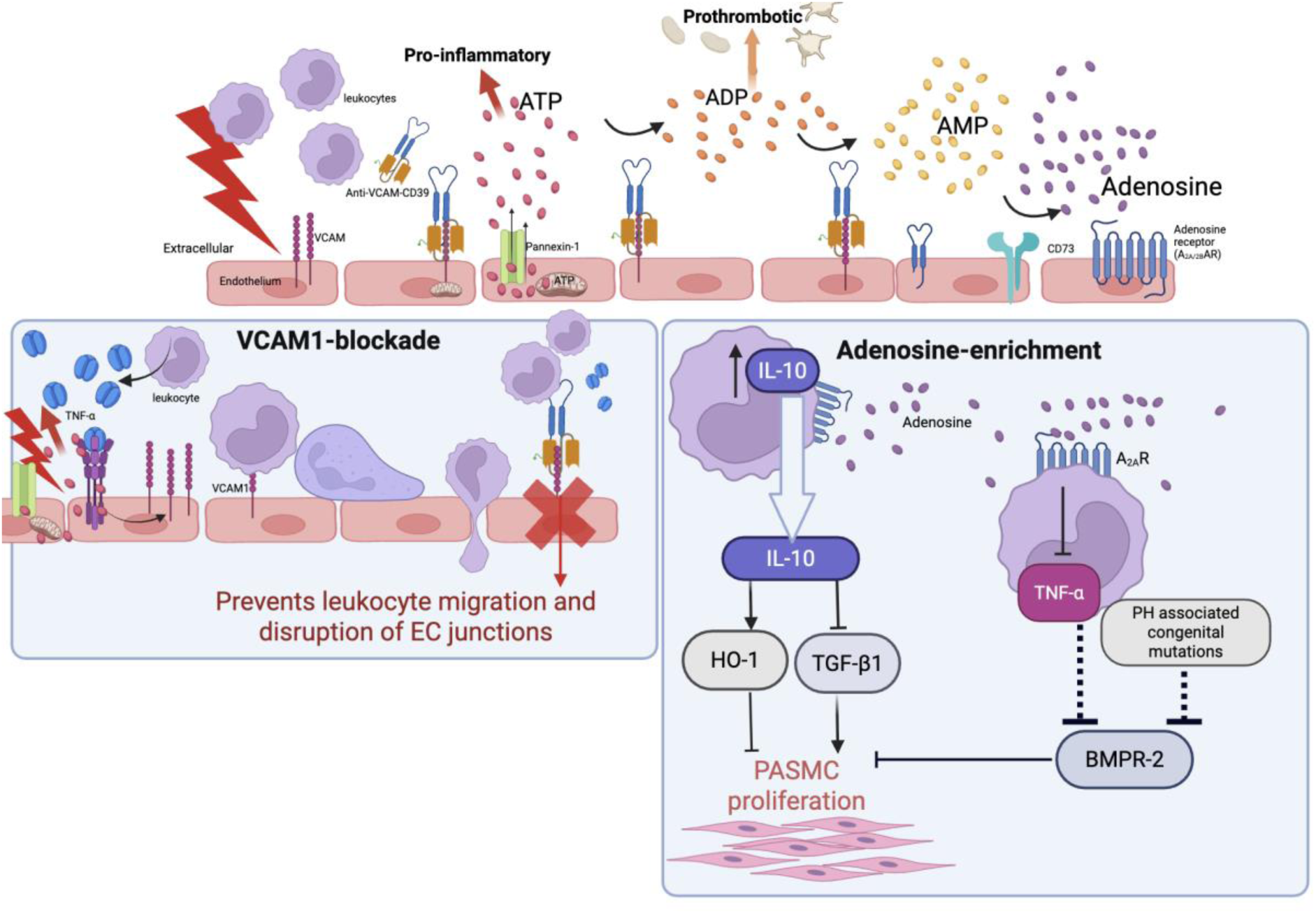
Schematic diagram of the hypothesized bifunctional mechanism at play with anti-VCAM-CD39 in MCTP-induced acute PAH Anti-VCAM-CD39 is a bifunctional molecule that blocks VCAM (left) and reduces pro-inflammatory ATP while increasing anti-inflammatory and antithrombotic adenosine in the vascular microenvironment (right). VCAM blockade reduces leukocyte adhesion and migration, likely preventing downstream actin remodeling and preserving EC junctions. CD39, working in concert with CD73, scavenges eATP and hydrolyzes it to adenosine. Adenosine promotes monocyte-derived IL-10 expression, which is an imperative regulator of PASMC proliferation via HO-1 signalling and TGF-β1 inhibition. TNF-⍺ disrupts BMPR-2 an potent modulator of pulmonary smooth muscle proliferation. The most common congenital forms of PH are due to mutations in BMPR-2. Adenosine inhibits TNF-⍺ which we hypothesized may restore BMPR2 signalling preventing PASMC proliferation.

Our MCTP-induced PAH model permits the delineation of contributions by VCAM-1 blockade and CD39 activity. Despite the known induction of VCAM-1 expression in PAH^11^, no prior reports have utilised VCAM-1 blockade alone in animal models of PH. VCAM-1 blockade by anti-VCAM alone protected animals from developing elevated RVSP at D10; however, it failed to prevent the right ventricular hypertrophy and small vessel remodeling characteristically seen in PAH. Clinically, right ventricular function emerges as a pivotal determinant of patients’ long-term survival ^70^ and late decompensatory ventricular dilatation can be seen in advanced PAH^71^ .Thus we hypothesize that VCAM-blockade adds to the benefit of CD39 for maximum benefit particularly if the dose of CD39 is to be kept at a level without the risk of bleeding.

A limitation of our study is that the development of PAH is not sustained after a single dose of MCTP. Multiple sequential challenges with MCTP may create a model with established chronic inflammatory signalling resembling human disease. However, the propensity of mice with CD39-deletion to develop PAH and the loss of endothelial expression of CD39 in human patients with PAH supports the use of MCTP to test the therapeutic benefit of CD39 supplementation. Whilst there are several treatments in clinical use for IPAH, there are clinical situations of abrupt decompensation^72^ . We propose that anti-VCAM-CD39 is a novel and safe strategy to reverse acute cardiorespiratory decompensation in PAH.

## Acknowledgments

We acknowledge Prof. Catriona McLean and Alfred Pathology for their assistance with patient lung samples. We also acknowledge the Monash Histology Platform, Monash University, for the provision of instrumentation, training and technical support.

## Authorship Contributions

AW: conceptualization, investigation, methodology, validation, data curation, and formal analysis, writing an original manuscript draft, manuscript review and editing

IS: data curation and formal analysis, investigation, methodology, validation, manuscript review and editing

NTL CS, IC, AV, VB, AW, YS, TW: investigation, methodology, data curation, validation

XW: methodology, investigation, manuscript review and editing

KHP: conceptualization, supervision, manuscript review and editing

SCR: conceptualization, ongoing collaboration, manuscript review and editing

HN: conceptualization, funding acquisition, investigation, methodology, project administration, resources, supervision, manuscript review and editing

MS: Conceptualization, data curation, formal analysis, funding acquisition, investigation, methodology, project administration, supervision, validation, manuscript review and editing

## Supplementary Material

### 1. Methods

#### 1.1. Design of the *anti-VCAM-CD39* construct expression and protein production

Anti-VCAM-1 ScFv was generated by Prof Claudia Gottstein and genetically fused to CD39. The construct was prepared in the baculovirus expression vector pBacPAK9 and contained the IL-3 signal sequence, FLAG epitope and cDNAs for scFv VCAM-1 and for soluble non-targeted human CD39.

Recombinant *anti-VCAM-CD39* was then produced using SF9 insect cells and purified by FLAG-Sepharose affinity-binding as detailed below.

SF21 insect cells in serum free media obtained from Walter and Eliza Hall Institute of Medical Research (WEHI) were thawed in 9mL of SF900 II SFM Insect cell media (Life Technologies, Australia) and centrifugated at 100xg for 5 minutes to remove DMSO. The cells were then resuspended in media at a cell density of 1x10^6^/mL and left to incubate on a shaker to ensure proper aeration of cells. The cells were incubated at 27°C for 2-3 days. The cells, at 1.5-2 x10^6^/mL final density, were then infected with the prepared anti-VCAM-CD39 virus at 10mL per L of cells and allowed to incubate with shaking for 48-72 hours.

The infected cells were placed on ice to reduce protein degradation and centrifugated at 100xg at 4°C. The supernatant was then transferred to 50mL tubes and centrifugated at 1000xg for 10minutes at 4°C. The supernatant was filtered and stored at -20°C till it was to be purified.

Anti-VCAM-CD39 secreted in the conditioned media was purified by immunoaffinity columns of anti-FLAG antibody (M2, Sigma-Aldrich Australia) covalently coupled to Sepharose and specifically bound anti-VCAM-CD39 was eluted via competitive binding with FLAG Peptide according to the manufacturer’s instructions. The FLAG-affinity purification column was stored at 50% glycerol in 1XTBS + 0.02% Sodium Azide. The column was allowed to drain and washed 3 times using 0.1M glycine in HCL, pH3.5. The column was then washed 5 times in 1XTBS to equilibrate the column for use. The stored supernatant is then prepared by neutralising the pH with 1M Tris (pH 8) and adding sodium chloride (NaCl) to a final concentration of 0.15M. The prepared supernatant was then loaded onto the FLAG column and allowed to run at 2.5mL/min flow rate. Peak fractions of protein were pooled and stored at -80°C.

#### 1.2. Production of control constructs, iScrVCAM-CD39 and ScFv-VCAM

To fully evaluate the efficacy of the bifunctional aspect of anti-VCAM-CD39, we used 2 control constructs. 1. Non-targeted VCAM-CD39, which contained a scrambled ScFv sequence but retained CD39 activity measured via adenosine activity assay 2. A single chain VCAM antibody, anti-VCAM.

Non-targeted CD39 (ScrVCAM-1-CD39) was produced in the same manner as above and similarly expressed in Sf21 cells followed by FLAG-affinity purification.

Anti-VCAM-1 was produced by Dr Aidan Walsh in Associate Professor Xiaowei Wang’s laboratory, Baker Heart and Diabetes Institute.Drosophila S2 insect cells were cultured in Express Five media (supplemented with 10 ml penicillin/streptomycin, and 90 ml L glutamine, all from ThermoFisher Scientific, USA) at 24 °C, 110 rpm, in a shaking incubator (Sheldon Manufacturing, USA). All procedures involving S2 cells were performed in sterile conditions, under laminar flow in a biological safety cabinet class II (Contamination Control Laboratories Pty Ltd, Australia), to avoid contamination. Cell counts were performed using a haemocytometer (LO–Laboroptik, UK) under a light microscope to calculate the concentration of S2 cells in culture. The cells were maintained at a concentration of between 1.5–3 × 106 to remain at an optimal concentration for growth.

Plasmid DNA was double filtered using 0.22 μm sterile filters and mixed with media and didodecyldimethylammonium bromide. This DNA mixture was allowed to incubate at RT for 20 minutes. Following this, the S2 cells were diluted to 1.5 × 106 cells per ml and the DNA mixture was added. The S2 cells were incubated at 24 °C, 110 rpm, for seven days as they produced antibodies.

After seven days, cells were removed from incubation and pelleted by ultracentrifugation at 6,000 × g for 30 minutes. The supernatant was collected, and the pellet was discarded. The supernatant was processed by Fast Protein Liquid Chromatography (FPLC), using an AKTA Pure system (GE Healthcare Life Sciences, United Kingdom). The supernatant passed through a polyhistidine affinity column (Qiagen, USA) and protein containing polyhistidine tags adhered to the column while media and impurities passed through. After washing the column with FPLC wash buffer, elution buffer containing imidazole at concentrations of 2%, 5%, and 10% v/v in wash buffer were successively passed through the column to remove weakly binding proteins. To collect protein, 100% elution buffer was passed through the column, which outcompeted polyhistidine binding to the column, freeing produced protein for collection in purified samples. Protein concentration within the fractions was measured at 280 nm and fractions containing high concentrations of protein were selected for dialysis.

Imidazole is not able to be used in animal studies; therefore it was removed following protein purification. The protein fractions were transferred to snakeskin dialysis bags (ThermoFisher Scientific, USA), and imidazole was dialysed into PBS at 4 °C.

#### 1.3. Echocardiogram

Animals were anaesthetised with isoflurane (1.5–3%). VisualSonics Vevo 2100 (VisualSonics Inc., Toronto, ON, Canada) and MS-550 D, 22–55 MHz. Pulse-wave doppler echo was used to record the pulmonary blood outflow at the level of the aortic valve in the short-axis view to measure the pulmonary acceleration time (PAT) and pulmonary ejection time (PET). The PAT/PET ratio is inversely correlated to right ventricular pressures (Thibault, 2010). The PAT and PET were taken as an average of 5–10 flow curves measured by an operator blinded to treatment groups using Vevo LAB Cardiac software (2023 FUJIFILM Visualsonics).

#### 1.4. Right Heart Catheterisation

Right heart cardiac catheterisation was performed by the Preclinical Cardiology Microsurgery and Imaging Platform at Baker Heart and Diabetes Institute. Mice were anaesthetised with isoflurane (2–4%). The right external jugular vein was isolated and a micro-tipped transducer catheter (a Millar pressure volume catheter (1.4Fr)) was inserted into the vein and advanced through the tricuspid valve and positioned in the RV for measurements of right ventricular pressure and contraction/relaxation rates. Ten or more pressure loops were measured by a blinded assessor to derive the mean RVSP for each animal.

#### 1.5. Western Blot

Whole lung tissue was homogenised (300 mg wet weight of tissue per 1 ml) of Lysis Buffer (TBS + 1% Triton X-100; Roche Australia). Protein concentration was quantified using BCA. Twenty microgram of total protein was subjected to SDS-PAGE (10% gel electrophoresed at 250 V for one hour). The proteins were transferred onto PVDF membranes (Merck Millipore, Australia) at a current of 150 mA for one hour, as previously described.^1,10^ VCAM-1 and Actin levels were detected by rabbit monoclonal anti-VCAM-1 (1:1000; 0.437 ng/ml; Abcam, UK) against a polyclonal swine anti-rabbit immunoglobulin (1:5000; Dako, Agilent USA) and by goat polyclonal anti-Actin (I-19)-HRP (1:500; 100 ng/ml; Santa Cruz, USA), respectively.

The Bio-Rad Chemidoc MP Imaging system (Bio-Rad Laboratories Pty. Ltd. Australia) was used to obtain chemiluminescence images. These images were quantitated by densitometry using Image Lab™ software (Bio-Rad Laboratories Pty. Ltd., Australia).

#### 1.6. eNOS ELISA

eNOS levels were determined using the commercial Mouse eNOS matched Antibody pair kit (Abcam, UK) according to the manufacturer’s instructions. Lung and liver lysates were prepared as described in sections 3.1.6.1 and 3.1.6.2 and thawed on ice on the day of testing. The provided capture antibody, diluted in coating buffer at 2 µg/mL (C3041-50CAP), was used to coat (50 µL per well) a 96-well flat bottom ELISA plate and incubated overnight at 4 °C. Plates were washed with wash buffer (1XPBS + 0.5% Tween-20) three times and then blocked with blocking buffer (1XPBS + 0.5% Tween-20 + 1%BSA; 100 µL per well) for two hours at room temperature to prevent non-specific binding. Lysates were diluted in standard diluent (1XPBS + 0.5% Tween-20 + 1%BSA) at a final concentration of 1:10. After washing the plates another three times, lysate samples and mouse eNOS standards (31.25 pg/mL to 2000 pg/mL) were added in duplicate at a final volume of 50 µL per well and incubated for two hours at room temperature on an orbital shaker. Plates were then washed three times with wash buffer, after which the detection antibody (0.5 µg/mL, 50 µL per well) was added to each well and incubated for one hour at room temperature on an orbital shaker. Plates were washed three times with wash buffer and HRP-conjugated Streptavidin at 0.05 µg/mL, 50 µL per well was added, after which the plate was incubated for one hour at room temperature on an orbital shaker. Plates were washed three times with wash buffer and 100 µL of TMB substrate solution was added to each well for 20 minutes of incubation at room temperature on an orbital shaker to allow colour to develop. After the development of colour was satisfactory, 0.18 M of H2SO4 was applied to stop the solution (50 µL per well) the absorbance (OD) was measured using the FLUOstar Optima Microplate Reader (BMG Labtech, Australia) at 450 nm. Total protein was quantitated using the BCATM protein assay. The amount of eNOS (pg) per microgram protein in each sample was calculated from the standard curve of the eNOS assay.

#### 1.7. Smooth muscle actin (SMA) Immunofluorescence

Frozen lung sections were thawed and protein blocked with BSA/Serum at 5% in PBS. Sections were then incubated with monoclonal mouse anti-human SMA (Dako, Clone 1A4, refM0851, lot 00053048, 1:500) followed by rabbit anti-mouse secondary antibody (Alexa Fluor 488, A11059, ThermoFisher, 1:1000). DAPI (Life Technologies, 50 ul per tissue section) was applied to sections followed by filtered Sudan black (0.3% in 70% EtOH). Images were captured using the Nikon TiE microscope or the Nikon A1R microscope (Nikon, Tokyo, Japan; Monash Micro Imaging Platform). Image quantification was performed using a novel script to measure blood vessel thickness by thresholding the SMA+ signal, which was developed to run in the public domain software Fiji (http://fiji.sc/Fiji). Quantification was performed by a blinded assessor. Ten or more vessels from each animal were quantified and the average taken.

#### 1.8. Immunofluorescent Staining

Paraffin fixed lung sections were acquired through Alfred Hospital Pathology with approval of the Alfred Hospital Ethics committee HREC 158/21. Lung tissue sections from patients with PAH were collected at the time of lung transplantation and a control cohort of patients who had had lung resection for alternative reasons who did not have PH were collected. CD31 (primary goat polyclonal antibody anti-CD31, R&D, AF3628, 1:200, secondary donkey anti-goat AF647, Invitrogen, A32849, 1:500) and CD39 (primary rabbit monoclonal anti-CD39, Abcam, ab223843, 1:200, secondary donkey anti-rabbit AF488, Jackson ImmunoResearch 711-545-152, 1:500) staining was performed by Monash Histology platform.

#### 1.9. Real-Time (RT)-PCR

Real-time Polymerase chain reaction (RT-PCR) was performed using the SensiFAST Lo-ROX Probe Kit (Bioline, Australia). Genes that were studied are described below, with gene expression assays purchased from ThermoFisher Scientific, Australia. RT-PCR was run on the QuantStudio 6 Real-Time PCR System (ThermoFisher Scientific, Australia) using the standard program. Genes analysed are listed in Table 3.2.

**Table 3.2:**
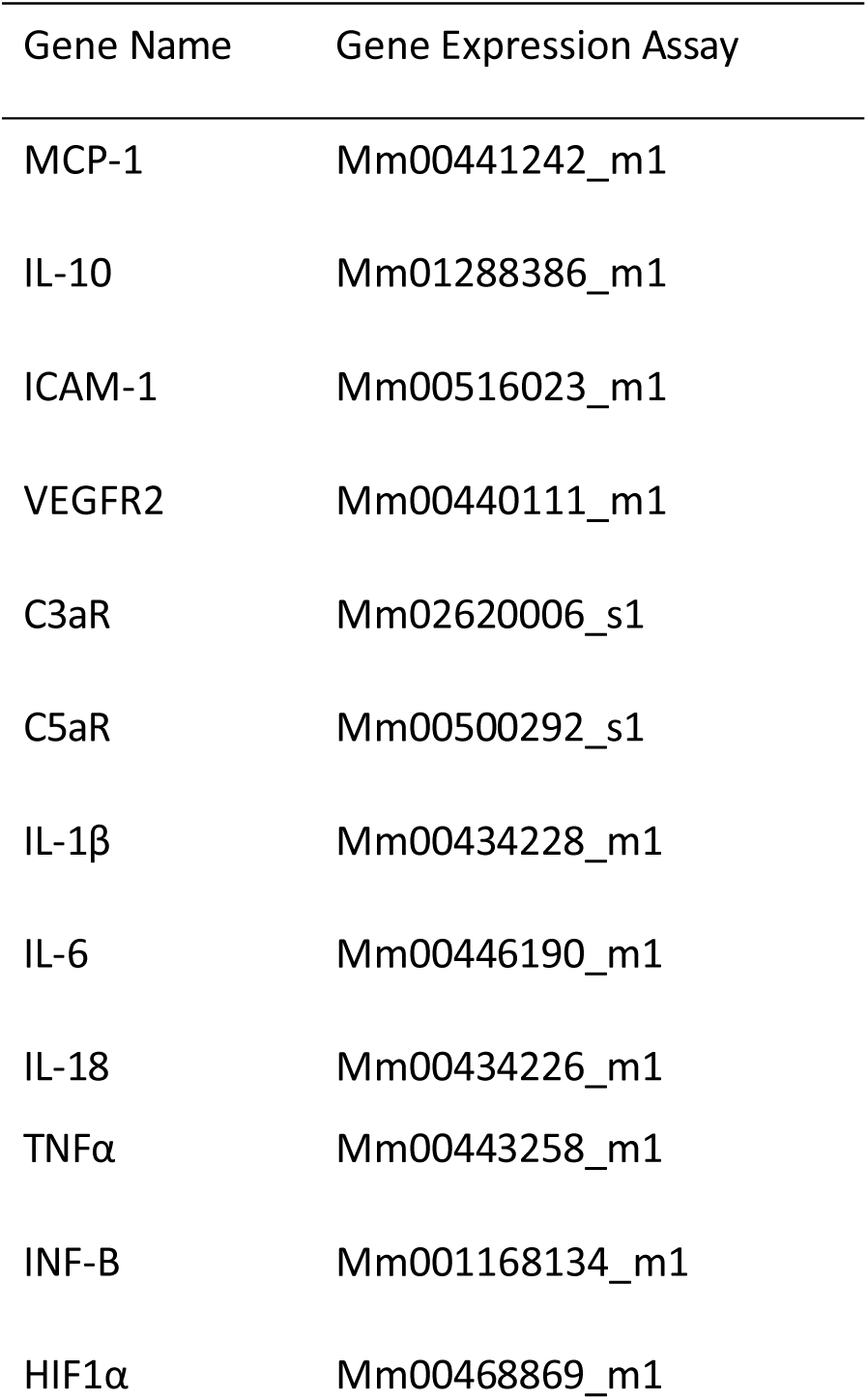
Gene expression assay.

### 2. Statistical analysis

This was performed using Prism 8 software (GraphPad, US). Comparison of experimental datasets was performed by one-way ANOVA with Uncorrected Fishers LSD (equal SD) or Brown-Forsythe and Welch ANOVA test with Dunnett’s T3 post-hoc analysis (unequal SD) for parametric data and Kruskal Wallis test for non-parametric data or two-way ANOVA with Uncorrected Fishers LSD as stated. In the absence of experimental values, this method gives the same P values and multiple comparisons tests as repeated measures ANOVA. P< 0.05 was considered significant. Outlier tests were also performed when appropriate. Differences between two groups were assessed by two-tailed student t-tests (unpaired or unpaired with Welch’s correction for parametric data and Mann-Whitney test for non-parametric data). P< 0.05 was considered significant.

### 3. Human Lung Tissue

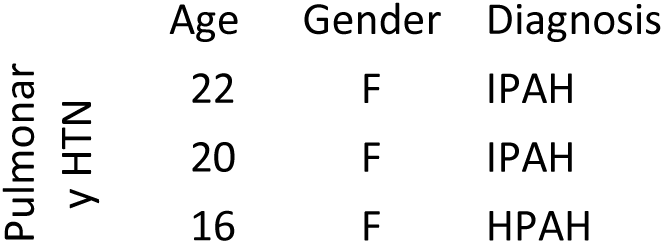
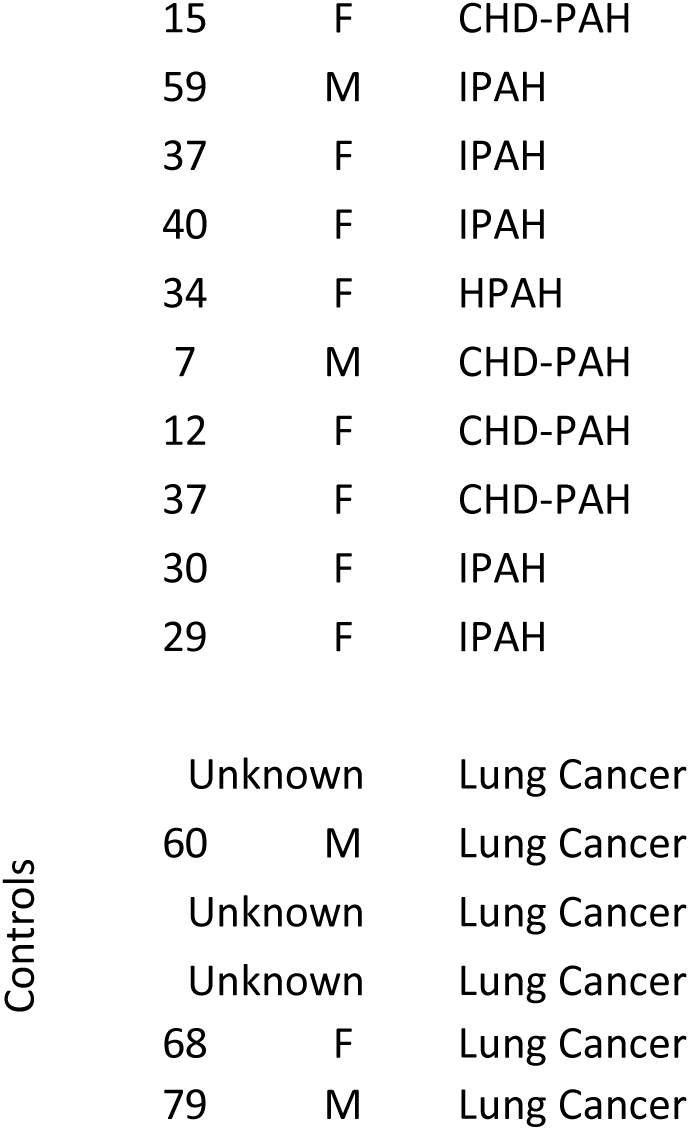

